# Finding MEMo: Minimum sets of elementary flux modes

**DOI:** 10.1101/705012

**Authors:** Annika Röhl, Alexander Bockmayr

**Affiliations:** Department of Mathematics and Computer Science, Freie Universität Berlin, Arnimallee 6, 14195 Berlin, Germany

## Abstract

Metabolic network reconstructions are widely used in computational systems biology for *in silico* studies of cellular metabolism. A common approach to analyse these models are *elementary flux modes* (EFMs), which correspond to minimal functional units in the network. Already for medium-sized networks, it is often impossible to compute the set of all EFMs, due to their huge number. From a practical point of view, this might also not be necessary because a subset of EFMs may already be sufficient to answer relevant biological questions. In this article, we study MEMos or minimum sets of EFMs that can generate all possible steady-state behaviours of a metabolic network. The number of EFMs in a MEMo may be by several orders of magnitude smaller than the total number of EFMs. Using MEMos, we can compute generating sets of EFMs in metabolic networks where the whole set of EFMs is too large to be enumerated.

## 1 Introduction

Genome-scale metabolic network reconstructions have been widely used in computational systems biology to develop *in silico* models of cellular metabolism [7, 8, 10, 30, 51]. Typically, a *metabolic network* 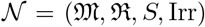 is given by a set 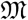 of (internal) metabolites, a set 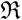 of reactions, a stoichiometric matrix 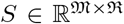, and a set 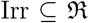 of irreversible reactions, see Fig. 1 for a small example. The set of reversible reactions is 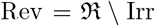. We always assume that the metabolic network is in steady-state, i.e., we analyse the (steady-state) *flux cone*

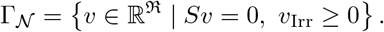

**Figure 1.**
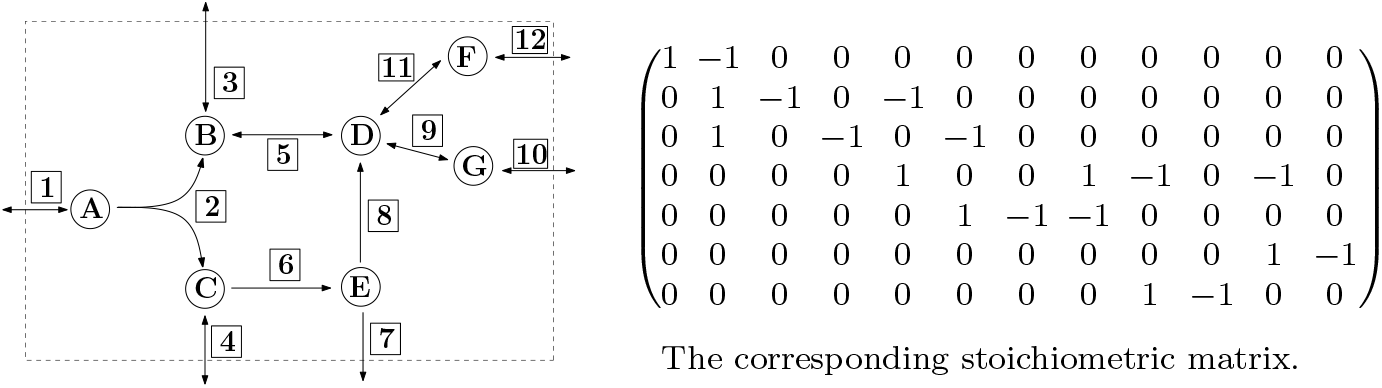
Example metabolic network with set of (internal) metabolites 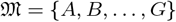, and set of reactions 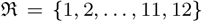, where Irr = {2, 6, 7, 8} is the set of irreversible reactions.

The elements 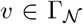 are called (feasible) *flux vectors* and can be interpreted as steady-state flux distributions over the network 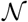.

The *support* of a flux vector 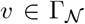 is the set 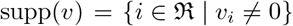 of active reactions in *v*. An *elementary flux mode* (EFM) [43] is a flux vector 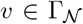, *v* ≠ 0 with inclusion-minimal support. One can show [44] that up to scaling an EFM is uniquely determined by its support. Therefore, two EFMs *e*, 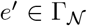 are considered to be the same if supp(*e*) = supp(*e*′). We denote by 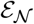 the finite set of EFMs in the metabolic network 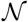. An EFM 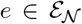 is called *reversible* if *e*, 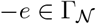, otherwise *e* is called *irreversible*. We denote the set of reversible resp. irreversible EFMs by 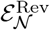 resp. 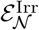.

### Example 1.1 (Elementary flux modes)

*In the network from Fig. 1 there are 18 EFMs with the following sets of active reactions:* {1, 2, 3, 4), {1, 2, 3, 5, 6, 8}, {1, 2, 3, 6, 7}, {1,2,3,6,8,9,10}, {1,2,3,6,8,11,12}, {1,2,4,5,9,10}, {1,2,4,5,11,12}, {1,2,5,6,7,9,10}, {1,2,5,6,7,11,12}, {1,2,5,6,8,9,10}, {1,2,5,6,8,11,12},{3,4,5,6,8},{3,5,9,10}, {3,5,11,12}, {4,6,7}, {4,6,8,9,10}, {4,6,8,11,12}, {9,10,11,12}. *The sets* {3,5,9,10}, {3,5,11,12}, {9,10,11,12} *are the supports of reversible EFMs, which can carry flux in both the forward and the backward direction.*

EFMs are a popular approach to analyse metabolic networks because every steady-state behaviour of the network can be represented with help of the EFMs [42, 43]. Formally speaking, the set 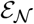 is a *generating set* or *conic basis* of the flux cone 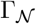. This means that every flux vector 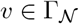 can be represented as *a conical combination* 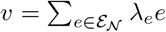, for some λ_*e*_ ≥ 0.

While the EFMs are of great theoretical and practical interest, already for medium-sized metabolic networks it is often not possible to enumerate all of them, since their number grows exponentially with the size of the network given metabolic network, see e.g. [6, 9, 12, 13, 32, 33, 35, 48, 52].

In this paper, we study inclusion-minimal sets of EFMs that generate the full flux cone 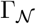. We study their mathematical properties and develop an algorithm to compute them. We start with a formal definition.

### Definition 1.2 (MEMo)

*Let* 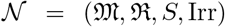 *be a metabolic network with flux cone* 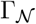 *and set of elementary flux modes* 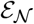. *A **M**inimum set of **E**lementary **Mo**eds or* MEMo *is an inclusion-minimal set* 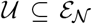 *such that every* 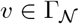 *can be represented as a linear combination*

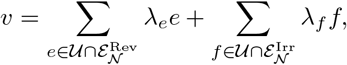

*for some* λ_*e*_, λ_*f*_ ∈ ℝ, *with* λ_*f*_ ≥ 0, *for all irreversible* 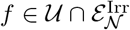.

A MEMo 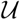 can be used to represent every other flux vector as a conical combination, in particular all EFMs not belonging to 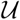. The minimum number of EFMs needed to represent the whole flux cone is in general much smaller than the number of all EFMs. Therefore, it may be possible to compute a MEMo in a reasonable amount of time even for large networks where the set of all EFMs can not be enumerated.

### 1.1 Intuition

We begin by providing some intuition on how to compute MEMos using the example in Fig. 1. Mathematically speaking, the flux cone 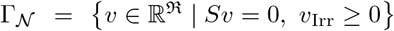 is a *finitely generated* polyhedral cone. This means that there exist finite sets 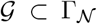 such that every flux vector 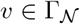 can be obtained as a conical combination 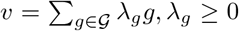 of the elements of 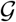. If the polyhedral cone is *pointed*, i.e., it does not contain any line {λ*v* | λ ∈ ℝ}, for some 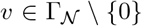, there exists a unique minimal set of generators. The elements of this set are the *extreme rays* (ERs) of the cone 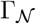, see Fig. 2 for illustration. In general, flux cones of metabolic networks are *non-pointed* because they contain *reversible flux vectors v* ≠ 0 for which both *v*, 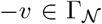. Non-pointed polyhedral cones do not have a unique minimal set of generators, see Fig. 3. The set of reversible flux vectors defines the *lineality space* 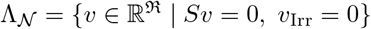. Following [26], a reversible reaction is called *fully reversible* if it can carry flux in a reversible flux vector. We denote the set of all fully reversible reactions in 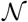 by Frev. The cone 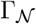 is pointed if and only if 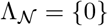.

**Figure 2.**
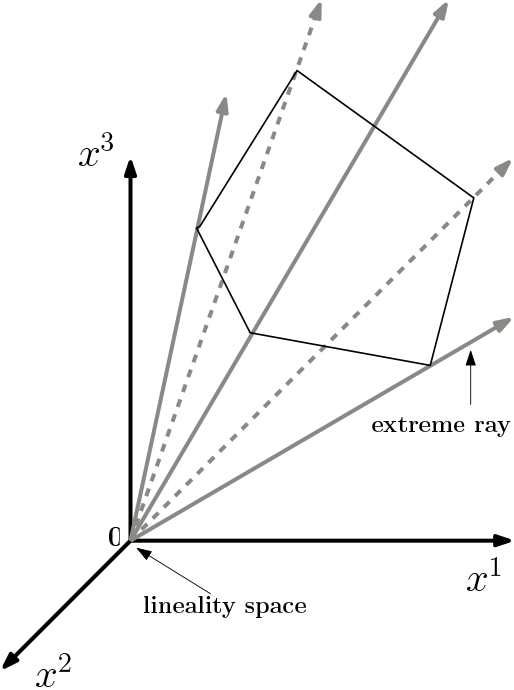
A pointed polyhedral cone in 3-dimensional space with 5 extreme rays. The lineality space is {0}.

**Figure 3.**
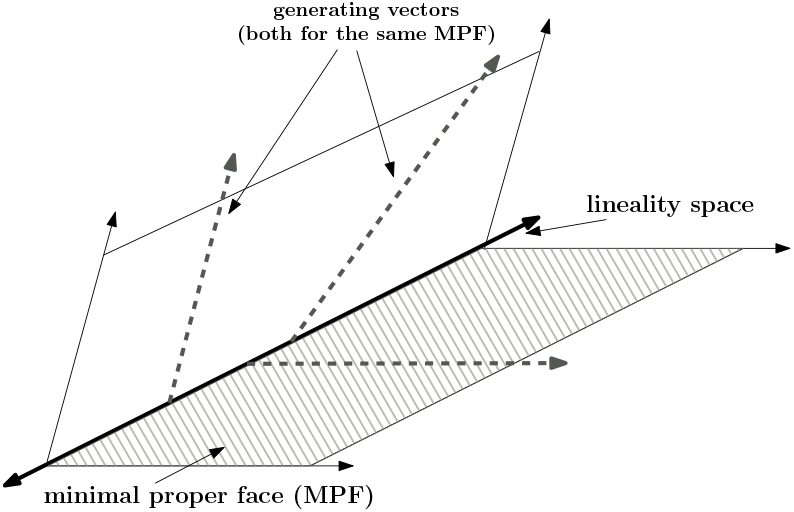
A non-pointed polyhedral cone in 3-dimensional space with a lineality space of dimension 1 and 2 minimal proper faces (MPFs) of dimension 2. The dashed vectors in one of the MPFs illustrate that the set of generating vectors is not unique: one of the two vectors can be used as well as any other vector in the MPF. To represent the whole polyhedral cone, one vector of each MPF is needed and one basis vector of the lineality space.

#### Example 1.3 (Reversible flux vectors and fully reversible reactions)

*The network in Fig. 1 contains flux vectors that consist of reversible reactions only. For example, v*^T^ = (0, 0, −1, 0, 1, 0, 0, 0, 1, 1, 0, 0) *is a vector with support* {3, 5, 9, 10} *and can operate in both directions. Thus, the reactions 3,5,9, and 10 are fully reversible and v*^T^ *is a reversible flux vector. In addition, reactions 11 and 12 of the network in Fig. 1 are fully reversible as well. On the other hand, reaction 4 is reversible, but not fully reversible: Whenever reaction 4 is active at least one irreversible reaction is active as well.*

As mentioned before, the set 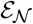 of EFMs is a finite generating set of the flux cone 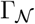. However, 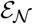 need not be minimal. A well-known method to compute the set of all EFMs consists in reducing the lineality space 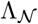 to {0} by splitting all reversible reactions in 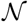, leading to a new network 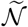. Splitting a reversible reaction *i* ∈ Rev means that reaction *i* is replaced by two reactions *i*^+^ and *i*^−^, where *i*^+^ represents the forward and *i*^−^ the backward direction of *i*.

#### Example 1.4 (Splitting all reversible reactions)

*In Fig. 4 all reversible reactions of the network in Fig. 1 are split. After splitting, there exists no reversible flux vector anymore. For example, a flux vector of the original network with support* {3, 5, 9, 10} *has now either the support* {3^−^, 5^+^, 9^+^, 10^+^} *or* {3^+^, 5^−^, 9^−^, 10^−^} *(the reverse direction).*

**Figure 4.**
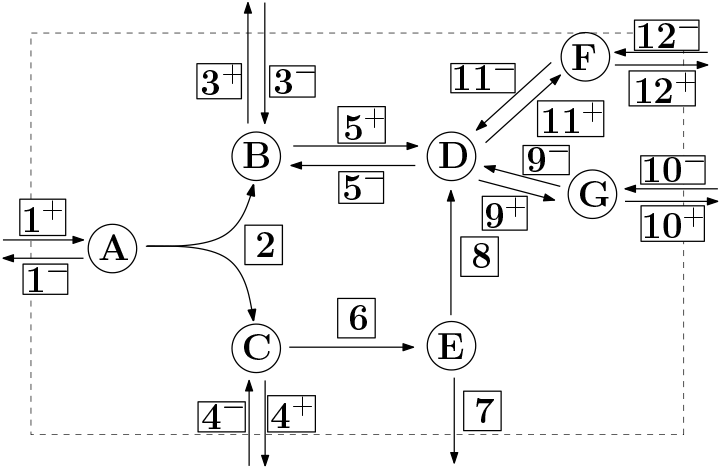
Same metabolic network as in Fig. 1, but with all reversible reactions split. For example, reaction 3 is split into 3^−^ and 3^+^.

The modified flux cone 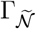, where all reversible reactions have been split, is pointed. Thus the extreme rays of 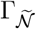 form a unique minimal set of generators. It can be shown that these extreme rays are in a 1-1 correspondence with the EFMs of the original network 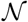 [5]. However, the set of 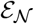 of EFMs need not be minimal for 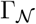.

#### Example 1.5 (Not all EFMs are needed)

*For the network in Fig. 1 the set of reactions* {1, 2, 3, 4} *is the support of the EFM v*^T^ = (1, 1, 1, 1, 0, 0, 0, 0, 0, 0). *The reactions* {3, 5, 11, 12} *are the support of an EFM as well w*^T^ = (0, 0, −1, 0, 1, 0, 0, 0, 0, 0, 1, 1). *Note that reaction 3 has a positive flux of 1 in v and a negative flux of* −*1 in w. The sum of v and w generates another EFM, u*^T^ = (1, 1, 0, 1, 1, 0, 0, 0, 0, 0, 1, 1), *consisting of the reactions* {1, 2, 4, 5, 11, 12}. *Thus, the EFM with the support* {1, 2, 4, 5, 11, 12} *is redundant and not required in a minimum generating set of EFMs for the network in Fig. 1.*

In this paper, we present a method for finding a minimum number of reversible reactions such that after splitting these reactions no feasible reversible flux vector exists in the modified network 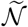. Therefore, the modified flux cone 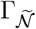 is pointed. We will show that the ERs of 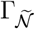 correspond to a MEMo of the original network 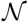, c.f. Def. 1.2.

#### Example 1.6 (Splitting a minimum set of reversible reactions)

*For the network in Fig. 1, it is sufficient to split only two reversible reactions, for example* 3 *and 9, see Fig. 5, to obtain a modified flux cone* 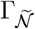 *which is pointed. There are 5 ERs of* 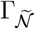, *which correspond to 5 EFMs in* 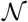. *Thus, only 5 instead of 18 EFMs are needed to generate* 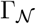. *The minimum set of reversible reactions to split is not unique. Instead of splitting* 3 *and* 9 *as in Fig. 5 one could also split the reactions* 10 *and* 12, *as in Fig. 6. Again, we obtain a MEMo containing 5 different EFMs, which however are different from the 5 EFMs in the MEMo obtained by splitting the reactions* 3 *and* 9. *Finally, not any set of two reversible reactions can be split to obtain a pointed cone. For example, splitting reactions* 11 *and* 12 *will not eliminate all reversible vectors, as shown in Fig. 7.*

**Figure 5.**
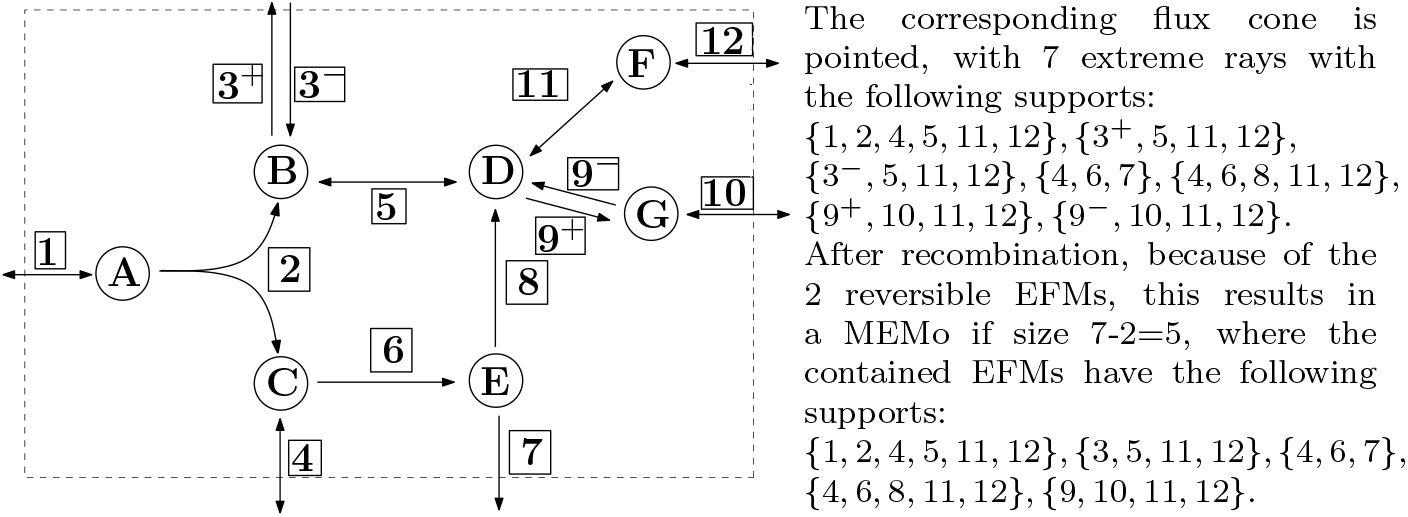
The fully reversible reactions 3 and 9 are split, leading to a pointed flux cone for the new network. The transformed network does not contain any reversible flux vectors.

**Figure 6.**
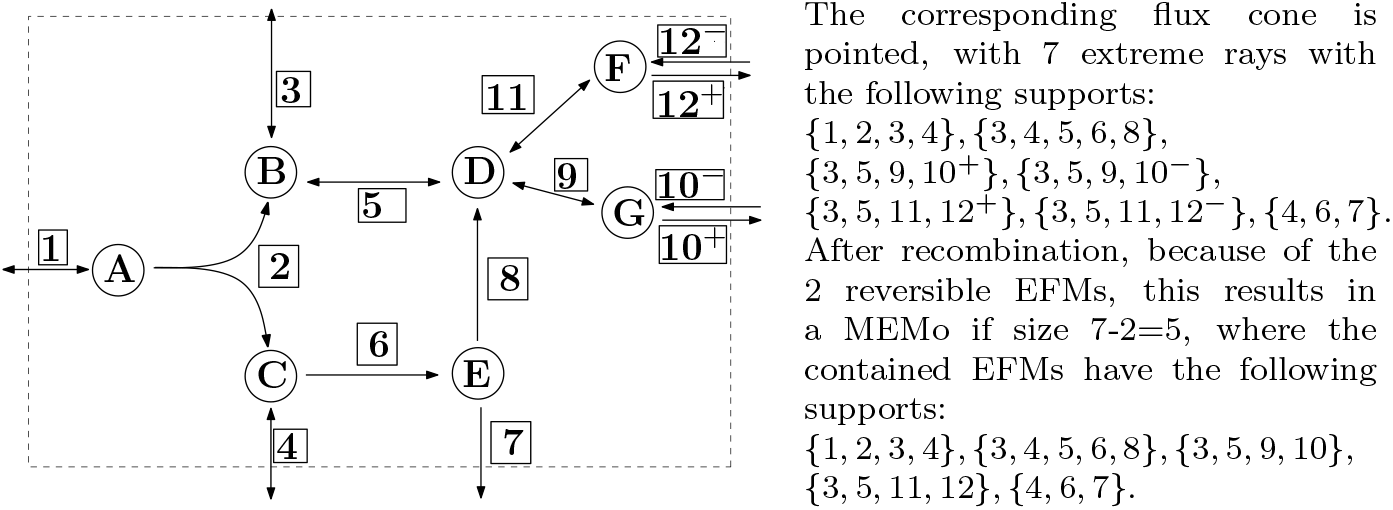
The fully reversible reactions 10 and 12 are split, leading to a pointed flux cone for the new network. The transformed network does not contain any reversible flux vectors.

**Figure 7.**
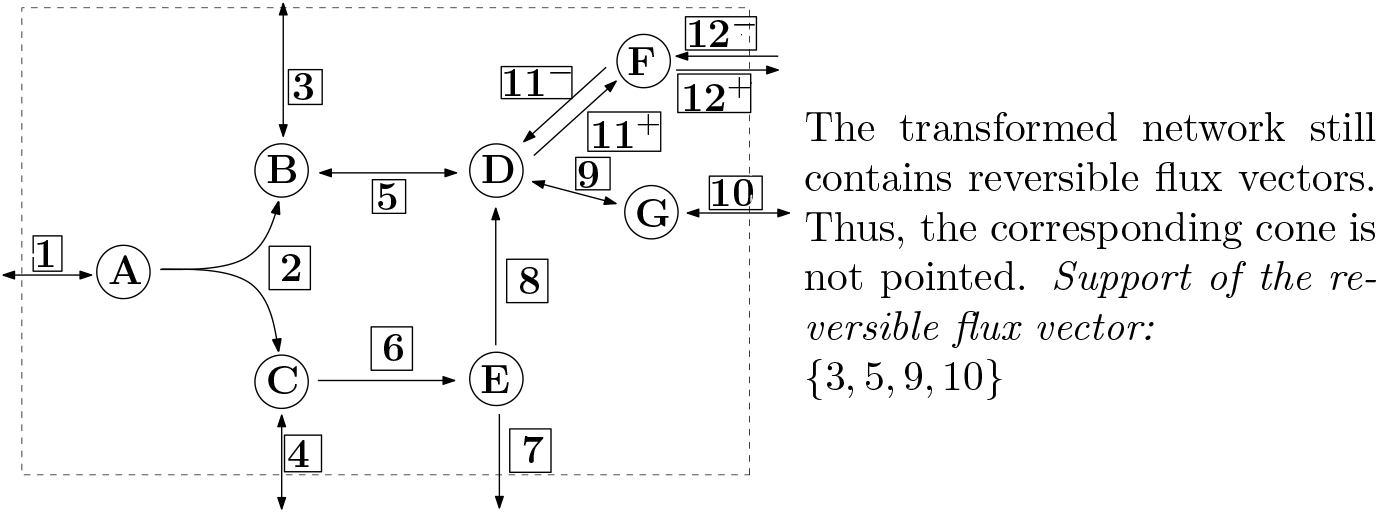
The fully reversible reactions 11 and 12 are split.

### 1.2 Contributions

The contributions of this paper are the following:

- We prove that whenever a (sub-)set of reversible reactions is split and the corresponding augmented flux cone is pointed, the extreme rays of the pointed augmented flux cone correspond to EFMs of the original network (Thm. 3.3).
- We introduce a method and provide an implementation for finding a minimum number of reversible reactions such that after splitting these reactions the resulting augmented flux cone is pointed. Additionally, we prove the correctness of the method (Thm. 4.2).
- We prove that the extreme rays of the pointed augmented flux cone (after splitting a minimum set of reversible reactions) correspond to a MEMo, needed to fully describe the underlying metabolic network (Thm. 5.5).
- We apply this method to several metabolic networks and the results show that the number of EFMs in a MEMo is by several orders of magnitude smaller than the number of all EFMs (Sect. 6).

The paper is organised as follows: We start in Sect. 2 with basic facts about polyhedral cones and their connection to metabolic networks. In Sect. 3, we focus on pointed flux cones and formally introduce splitting of reversible reactions. We prove that if splitting results in a pointed polyhedral cone, then the extreme rays of the augmented flux cone correspond to EFMs of the original metabolic network. In Sect. 4 we show how to find a minimum set of reversible reactions to split such that the resulting augmented flux cone is pointed. In Sect. 5 we prove that the extreme rays of the pointed augmented flux cone (after splitting a minimum set of reversible reactions) correspond to a MEMo. Finally, in Sect. 6 we discuss and provide computational results for various genome-scale metabolic networks that were obtained by the software that we implemented. Related and future work are discussed in Sect. 7 and Sect. 8.

A preliminary version of this article appeared as a conference paper in [37].

## 2 Polyhedral cones

After providing the intuition underlying MEMos, we now formally introduce the method. For this, we start with some mathematical background on polyhedral cones, see [41] for more details.

### 2.1 Notation

For a vector *x* ∈ ℝ^*n*^ we denote by *x*_*i*_ the *i*-th element of *x*. For a set *H* of indices, *x*_*H*_ is the subvector of *x* corresponding to these indices. We use superscripts to refer to different vectors. For example, {*x*^1^, …, *x*^*t*^} denotes a set of *t* vectors and we use a set of indices *I* here as well: *x*^*I*^ = {*x*^*i*^ | *i* ∈ *I*}. If all elements of a vector should be greater or equal to zero we write *x* ≥ 0 instead of *x*_*i*_ ≥ 0 for all *i*. With 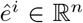 we denote the *i*-th unit vector. We denote by *A*_*i*,∗_ the *i*-th row and by *A*_∗,*j*_ the *j*-th column of the matrix *A*. With *A*_*H*,∗_ resp. *A*_∗,*H*_, we refer to a set of rows, resp. to a set of columns of *A*. We denote the rank of a matrix *A* ∈ ℝ^*m*×*n*^ with *ρ*(*A*), where *ρ*(*A*) is the maximum number of linearly independent columns in *A*. The horizontal resp. vertical concatenation of matrices *A, B* is denoted by 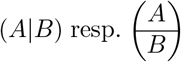. Finally **0**_*m,n*_ are zero matrices of the size *m* × *n*.

### 2.2 Polyhedral cones

A subset Γ ⊆ ℝ^*n*^ is called a *cone* if any *conical combination* of two elements *x, y* ∈ Γ belongs to Γ again, i.e., λ*x* + *μy* ∈ Γ, for any non-negative λ, *μ* ∈ ℝ_≥0_. An element *x* ∈ Γ \ {0} is called a *ray* of Γ. A cone Γ is called *polyhedral* if there exists a matrix *A* ∈ ℝ*m*×*n* such that

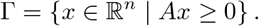

A cone Γ is *finitely generated* if there exists a finite set of generators {*g*^1^, …, *g*^*t*^} ⊆ Γ such that every element *x* ∈ Γ can be written in the form 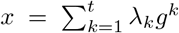, for some non-negative λ_*k*_ ∈ ℝ_≥0_. By a classical theorem of Farkas-Minkowski-Weyl, a cone is polyhedral if and only if it is finitely generated. The *dimension* dim(Γ) of a polyhedral cone Γ is the maximum number of affinely independent points of Γ minus 1. For a polyhedral cone Γ = {*x* ∈ ℝ^*n*^ | *Ax* ≥ 0}, the linear subspace

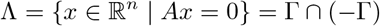

is called the *lineality space* of Γ. An inequality *a*^T^*x* ≥ 0, *a* ∈ ℝ^*n*^ \ {0} is *valid* for Γ if Γ ⊆ {*x* ∈ ℝ^*n*^ | *a*^T^*x* ≥ 0}. A set *F* ⊆ Γ is called a *face* of Γ, if there exist a valid inequality *a*^T^*x* ≥ 0 for Γ such that *F* = Γ ⋂ {*x* ∈ ℝ^*n*^ | *a*^T^*x* = 0}. *F* is called a *facet* if dim(*F*) = dim(Γ) − 1. A face *F* is called a *minimal proper face* (MPF) if dim(*F*) = *t* + 1, where *t* is the dimension of the lineality space Λ, see Fig. 3 for illustration.

A polyhedral cone Γ is called *pointed* if Λ = {0}. This means that whenever *x* ∈ Γ, *x* ≠ 0, we have −*x* ∉ Γ. In other words, Γ does not contain any line {λ*x* | λ ∈ ℝ}, for *x* ≠ 0. By basic linear algebra, the cone Γ is pointed if and only if the matrix *A* has full column rank, i.e., *ρ*(*A*) = *n*. Two rays *x, x*′ ∈ Γ are considered identical if *x*′ = λ*x* for some λ > 0. A ray *x* ∈ Γ \ {0} is called an *extreme ray* (ER) of Γ, if there exist no two linearly independent rays *x*^1^, *x*^2^ ∈ Γ such that *x* = *x*^1^ + *x*^2^. If Γ is pointed, the minimal proper faces have dimension one. They form a unique minimum set of generators {*g*^1^, …, *g*^l^} ⊆ Γ which correspond to the ERs of Γ, see Fig. 2 for illustration.

For a more algebraic characterisation of the ERs of a pointed polyhedral cone we can use the *zero set Z*(*x*) = {*i* | *x*_*i*_ = 0}, which denotes the set of indices of entries of a given vector *x* which are zero. According to [12, 19], a ray *x* of a pointed polyhedral cone Γ = {*x* ∈ ℝ^*n*^ | *Ax* ≥ 0} is an ER if and only if *ρ*(*A*_*Z*(*Ax*),∗_) = *n* − 1, where *ρ*(*A*) = *n*.

The set of feasible flux vectors in a metabolic network 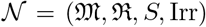 defines a polyhedral cone

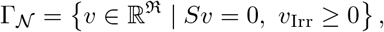

which is called the *flux cone*. As before, we denote by 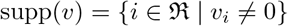 the *support of* 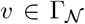 and by supp_Irr_(*v*) = {*i* ∈ Irr | *v*_*i*_ ≠ 0} the *support of v in* Irr. An *elementary flux mode* in 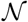 is a flux vector 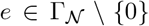 with inclusion-minimal support. According to the *rank test* [19, 52], a flux vector 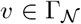 is an EFM if and only if *ρ*(*S*_∗,supp(*v*)_) = |supp(*v*)| − 1, where |supp(*v*)| is the number of non-zero entries of *v*.

[26] introduced a notion similar to the EFMs, using the set of irreversible reactions Irr. A *minimal metabolic behavior* is an inclusion-minimal non-empty set *D* ⊆ Irr for which there exists 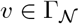 such that suppIrr(*v*) = *D*.

#### Example 2.1 (Minimal metabolic behaviours)

*The network in Fig. 1 has three MMBs, namely* {2}, {6, 7}, *and* {6, 8}. *The minimal metabolic behaviour* {6, 8} *is present in 8 EFMs with the supports* {1, 2, 3, 5, 6, 8}, {1, 2, 3, 6, 8, 9, 10}, {1, 2, 3, 6, 8, 11, 12}, {1, 2, 5, 6, 8, 9, 10}, {1, 2, 5, 6, 8, 11, 12}, {3, 4, 5, 6, 8}, {4, 6, 8, 9, 10}, {4, 6, 8, 11, 12}. *The last 3 EFMs, for which* supp_Irr_(·) = {6, 8}, *are all contained in the minimal proper face associated with the MMB* {6, 8}, *cf. [26]. The remaining 5 EFMs, for which* supp_Irr_(·) = {2, 6, 8}, *lie in the interior of the cone* 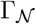.

The following proposition summarises what is known about the relationship between extreme rays, EFMs, and MMBs.

#### Proposition 2.2

*Let* 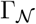 *be the flux cone of a metabolic network* 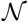.

1. *If all reactions are irreversible, then* 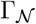 *is pointed and the extreme rays of* 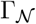 *are exactly the EFMs of* 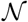 *[13, 43]*.
2. *If there are reversible reactions and* 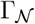 *is pointed, then the extreme rays of* 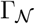 *form a subset of the set of EFMs in* 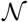. *However, not every EFM needs to be an extreme ray [19]*. *The extreme rays are exactly the vectors in* 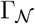 *with inclusion-minimal support in* Irr.
3. *If* 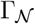 *is not pointed, then there is a 1-1 correspondence between the MMBs of* 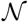 *and the minimal proper faces of* 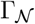 *[26]*.

**Proof:** We have only to prove the last part of 2). By [26] (or using 3)) there is a 1-1 correspondence between the minimal proper faces of a flux cone and the MMBs. All vectors in a minimal proper face *F* have the same support in Irr, which is exactly the MMB corresponding to *F* . If a polyhedral cone is pointed, the minimal proper faces are the extreme rays.

In general, a metabolic network may contain reversible reactions together with reversible flux vectors *v* ≠ 0, for which *v*, 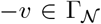, and the flux cone is non-pointed. For example, in Fig. 1, there is a reversible flux vector with the support {3, 5, 9, 10}. If there exist reversible flux vectors, a minimal generating set of the flux cone is not unique anymore. It may consist of an arbitrary flux vector for each minimal proper face of the flux cone, together with a vector space basis of the lineality space, see [41, Sect. 8.8]. Computing a minimal generating set by one of the standard software tools, usually does not deliver a set of EFMs. In order to obtain a minimal generating set consisting of EFMs, one can transform the flux cone into a pointed cone. In the next section, we explain how this can be done.

## 3 Splitting reversible reactions

In many cases it is desirable to have a cone Γ which is pointed. If *x* ≥ 0 for all *x* ∈ Γ, then the resulting cone will be pointed. In general, however, a variable *x*_*i*_ ∈ ℝ can take negative values and the constraint *x*_*i*_ ≥ 0 does not hold. To overcome this problem, a well-known method also used in linear programming is to *split* variables. Splitting a variable *x*_*i*_ ∈ ℝ means replacing it by two non-negative variables 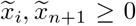, such that 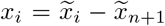. Note that this will change the structure of the cone and increase the dimension of the underlying vector space by 1. To describe this transformation formally and for several variables, we use a map π_*I*_ : ℝ^*n*^ → ℝ^*n*+|*I*|^, where *I* = {*i*_1_, …, *i*_*s*_} denotes the set of variables 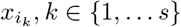 to be split. For *x* ∈ ℝ^*n*^ we get 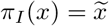 with 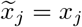, for all *j* ∈ {1, …, *n*} \ *I*, and for each *i*_*k*_ ∈ *I*:

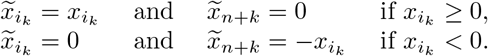

Using the map π_*I*_, a polyhedral cone Γ = {*x* ∈ ℝ^*n*^ | *Ax* ≥ 0}, with *A* ∈ ℝ^*m*×*n*^, is mapped to the *augmented polyhedral cone*

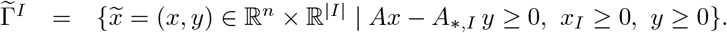

The inverse transformation 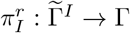 maps each vector 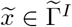 to 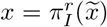 such that 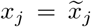, for all *j* ∈ {1, …, *n*} \ *I*, and 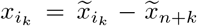, for all *i*_*k*_ ∈ *I, k* ∈ {1, …, *s*}. If we apply 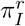, we say that we *recombine* the variables that were split before.

We now turn to the lineality space of 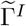. Generalising a result from [25], we get:

### Proposition 3.1

*Let* Γ ⊆ ℝ^*n*^ *be a polyhedral cone with lineality space* Λ. *For a set of variables I* ⊆ {1, …, *n*}, *the lineality space* 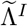 *of the augmented polyhedral cone* 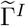 *is given by*

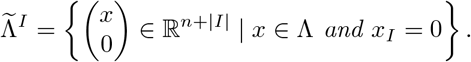

**Proof:** For the cone Γ = {*x* ∈ ℝ^*n*^ | *Ax* ≥ 0} the lineality space is Λ = {*x* ∈ ℝ^*n*^ | *Ax* = 0}. Splitting the variables in *I* delivers the cone 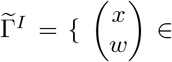 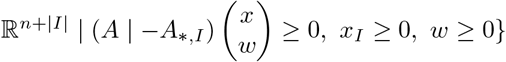 For the lineality space we get:

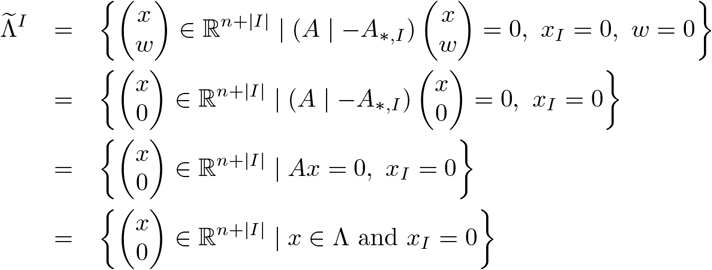

### Corollary 3.2

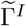 *is pointed if and only if ρ*(*A*_∗,{1,…,*n*}\*I*_) = *n* − |*I*|.

In the case of a metabolicxs network 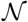, the variables corresponding to the irreversible reactions can take only non-negative values. In order to obtain a pointed cone, one may split all reversible reactions into two irreversible ones, see Fig. 4 and Example 1.4. This leads to the pointed augmented ux cone 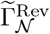. The uniquely determined extreme rays of 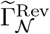 are called *extreme currents* [5]. It has been shown in [13] that after recombination and up to so-called 2-cycles, the extreme currents correspond exactly to the EFMs of the metabolic network 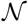. For each split reaction *i*_*k*_ ∈ Rev, the corresponding *2-cycle* is the extreme ray 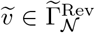 with 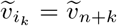, and *v*_*i*_ = 0 otherwise. These cycles do not have a biological meaning and can be eliminated (they become zero after applying 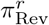).

Splitting all reversible reactions may highly increase the number of variables and thus the dimension of the vector space where the augmented flux cone lives in. Already for medium-sized networks, the number of extreme rays of 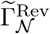, which corresponds to the number of EFMs (up to the 2-cycles) of 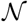, will be huge and therefore computing the whole set may not be feasible or desirable. Already in [40], it has been shown that splitting only the *internal* reversible reactions delivers an augmented flux cone which is pointed (assuming that there is at most one exchange reaction per internal metabolite). After recombination, the extreme rays of this cone are called *extreme pathways* (EPs). However, the number of EPs can be very large and usually the set of all EPs cannot be computed for genome-scale metabolic networks, see Sect. 7.1 for further discussion.

Extending the results in [13], we show next that if splitting a subset *I* ⊆ Rev of reversible reactions results in a pointed augmented flux cone 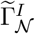 then the extreme rays of 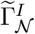 after reconfiguration define a subset of EFMs in the metabolic network 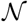.

### Theorem 3.3

*Let* 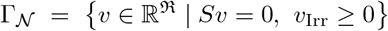 *be the flux cone of a metabolic network* 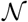. *Let I* ⊆ Rev *be a set of reversible reactions such that the augmented cone* 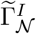 *obtained by splitting the reactions in I is pointed. With exception of the* 2*-cycles, the extreme rays of* 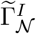 *after recombination are elementary flux modes in* 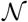.

**Proof:** By splitting the reactions in *I* we get the metabolic network 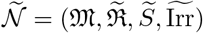 with 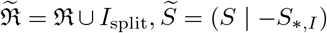 and 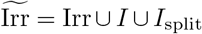. Here, *I*_split_ is the set of additional irreversible reactions obtained by splitting the reactions in *I*. The corresponding flux cone is

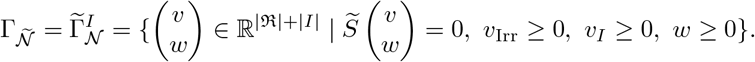

Let 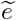 being an ER of 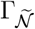 and 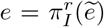 the corresponding flux vector in 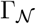. By Prop. 2.2, the ERs of the pointed cone 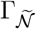 are EFMs in 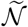. By the rank test for EFMs, this implies 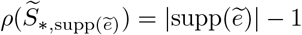.

Suppose first that for all *i*_*k*_ ∈ *I* we have 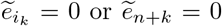, i.e., there is no non-zero flux through both directions of a former reversible reaction. Then 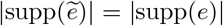 and 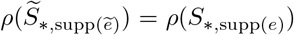 (by the definition of 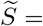 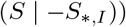). It follows that *ρ*(*S*_∗,supp(*e*)_)) = |supp(*e*)| − 1. Using again the rank test, this shows that e is an EFM in 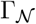.

Suppose now that there exists *i*_*k*_ ∈ *I* with 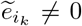 and 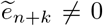. Since 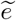 is an EFM of 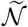, it follows 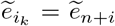 and 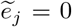, for all 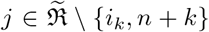. Hence, 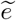 is a 2-cycle, which we excluded from our considerations.

Thus, whenever the augmented flux cone 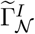 is pointed, the recombined extreme rays of 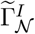 correspond to a subset of EFMs in the original metabolic network 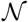. The remaining EFMs of 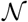 correspond to rays inside of 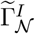 and can be obtained from the extreme rays by conical combinations. In the following, we develop a method for determining a minimum set of reversible reactions that have to be split in order to obtain a pointed cone.

## 4 Splitting a minimum number of reversible reactions

We show in this section that there exist minimum sets of reversible reactions such that, after splitting these reactions, the augmented flux cone becomes pointed. These sets always contain *t* fully reversible reactions, where *t* is the dimension of the lineality space. We note that splitting one reaction reduces the dimension of the lineality space by at most 1 [25]. Therefore at least *t* reversible reactions have to be split in order to obtain a pointed cone.

### Definition 4.1 (minFrev)

*Let* 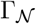 *be the flux cone of a metabolic network* 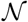. *Let t be the dimension of the lineality space* 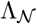. *A* minimum set of fully reversible reactions *is a set* minFrev *of t fully reversible reactions such that the flux cone* 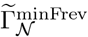 *obtained by splitting the reactions in* minFrev *is pointed. We denote by* minFrev_split_ *the set of the t additional irreversible reactions obtained by splitting. Thus*

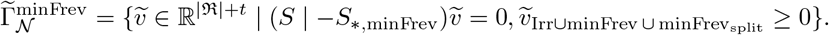

The next theorem states that for any metabolic network 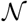 there exists a minimum set of fully reversible reactions minFrev in the sense of Def. 4.1. To find such a set, we use a well-known concept from matrix theory [14, p.85]. A matrix *B*^rcef^ ∈ ℝ^*m*×*n*^ is in *reduced column echelon form* if the following properties hold (see Fig. 8):

1. The first non-zero element in column *k* is a 1 in row *j*_*k*_, for *k* = 1, 2, …, *r* (this 1 is called a *pivot*).
2. 1 ≤ *j*_1_ < *j*_2_ < ⋯ < *j*_r_ ≤ *m* (i.e., for each change in columns from left to right, the pivot appears in a lower row).
3. For *k* = 1, …, *r*, the pivot in column *k* is the only non-zero element in row *j*_*k*_.
4. Each of the last *n* − *r* columns consists entirely of zeros.

**Figure 8.**
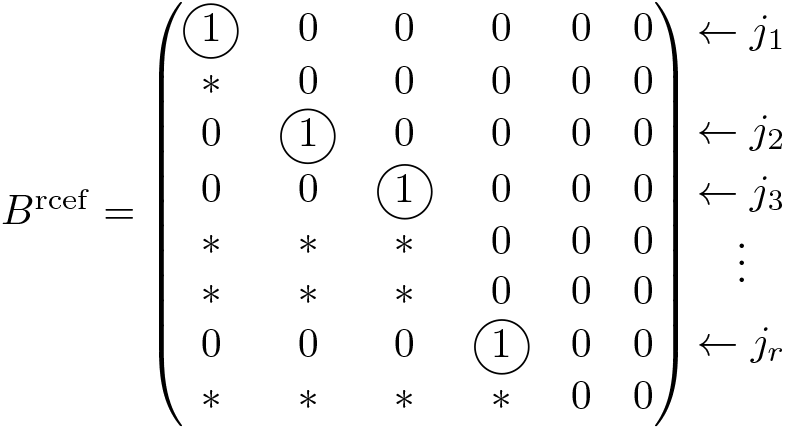
*B*^rcef^ is a matrix in reduced column echelon form, where ∗ are appropriate values in ℝ.

### Theorem 4.2

*Let* 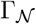 *be the flux cone of a metabolic network* 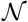. *If t is the dimension of the lineality space* 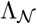, *then there exists a set* minFrev *of t fully reversible reactions such that the cone* 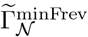 *obtained by splitting the reactions in* minFrev *is pointed.*

**Proof:** The idea of the proof is the following. As a vector space, the lineality space 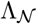 has a basis consisting of *t* linearly independent vectors. In order to obtain a pointed flux cone, we intuitively have to destroy 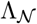, i.e., all vectors in the basis. We do this by splitting one reaction in each vector of the basis. To make sure that each of these reactions is present in exactly one basis vector, we make use of the reduced column echelon form, cf. Fig. 8.

We may assume that 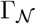 is not pointed, hence 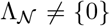. Let 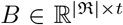 be a matrix whose columns 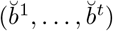 form a basis of the lineality space 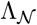, where *t* ≥ 1 is the dimension of 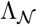. By applying elementary column operations, we can obtain the reduced column echelon form *B*^rcef^ of *B*, see Fig. 8, which is uniquely determined by *B* [14, p.85]. The columns (*b*^1^, …, *b*^*t*^) of *B*^rcef^ define again a basis of 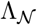. The row indices *j*_1_, …, *j*_r_ for the pivot 1’s in *B*^rcef^ are exactly the indices of the fully reversible reactions we are looking for. To see this, define minFrev = {*j*_1_, …, *j*_*r*_}. From *t* = *ρ*(*B*) = *ρ*(*B*^rcef^) = *r*, we get *r* = *t*. Since 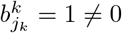, for *k* = 1, …, *r* = *t*, we have minFrev ⊆ Frev.

After splitting the reactions in minFrev we get the augmented flux cone 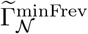, which by Prop. 3.1 has the lineality space

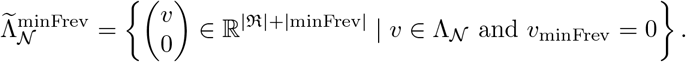

Since (*b*^1^, …, *b*^*t*^) defines a basis of 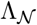, any 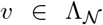 can be written as 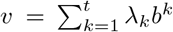, for some λ_*k*_ ∈ ℝ. The matrix *B*^rcef^ is in reduced column-echelon form. This means that for each *j* ∈ minFrev there is exactly one 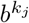 with 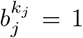, and for all *k* ≠ *k*_*j*_, it holds 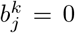. Since for 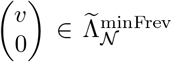 we have 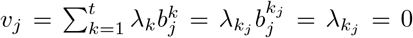, for all *j* ∈ minFrev, it follows λ_*k*_ = 0, for all *k* ∈ {1, …, *t*}. This implies *v* = 0 and thus 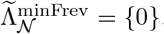, 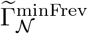 which proves that 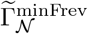 is pointed.

Based on the previous proof, Algorithm 1 summarizes how to find a minimum set minFrev such that 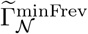 is pointed. First we compute a basis of the lineality space 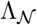 of 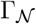. Let *B* be the matrix whose columns are the vectors of this basis. Next we transform *B* in the reduced column echelon form (e.g. in `MATLAB` using `rref`). The row indices of the pivots then correspond to the indices of the reactions that form the set minFrev.

### Algorithm 1: Finding a minimum set of fully reversible reactions

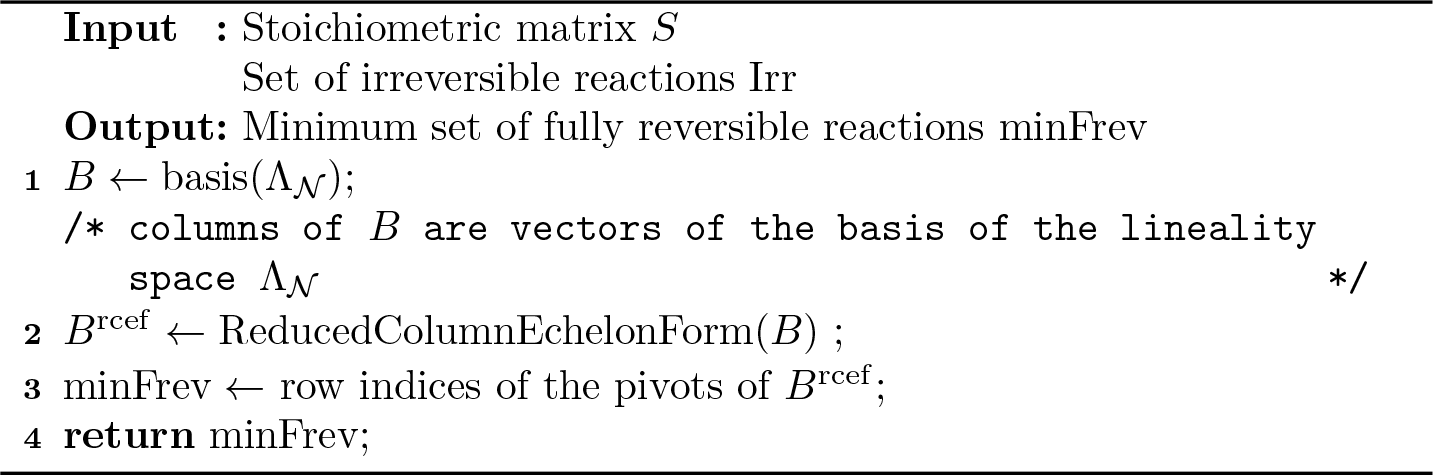

We note that the set minFrev computed by Algorithm 1 does not depend on the particular basis *B* of 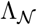 because two matrices *B, B*′ with the same column space have the same reduced column-echelon form [14].

### Example 4.3 (Compute minFrev)

*The flux cone for the network in Fig. 1 has a lineality space of dimension 2. Therefore any set* minFrev *contains two reactions. The lineality space can be described by the matrix*

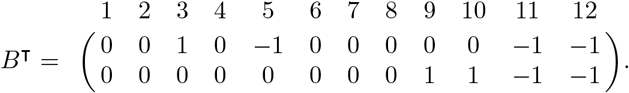

*We can see that only the fully reversible reactions* 3, 5, 9, 10, 11 *and 12 are active in the lineality space. Furthermore, B is already in reduced column echelon form and thus B* = *B*^rcef^. *The pivot elements are 3 and 9, therefore* minFrev = {3, 9}. *These are the reactions which are split in Example 1.6. Splitting reactions 3 and 9 delivers an augmented flux cone which is pointed. This can be seen from the modified network in Fig. 5, since there exists no feasible flux vector that is reversible.*

## 5 MEMo: Minimum set of Elementary Modes

The main result of this section is that the extreme rays of an augmented flux cone 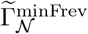, where minFrev is a minimum set of fully reversible reactions, after recombination form a Minimum set of Elementary Modes or MEMo, cf. Def. 1.2. We start by using Prop. 2.2 to prove the following result.

### Proposition 5.1

*Let* minFrev *be a minimum set of fully reversible reactions in a metabolic network* 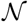 *such that* 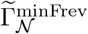 *is pointed. Then the set of extreme rays of the augmented cone* 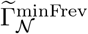 *does not contain any 2-cycle.*

**Proof:** By the definition of the lineality space 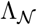 and the set Frev, we have 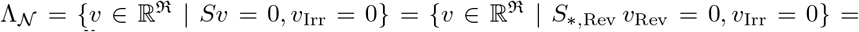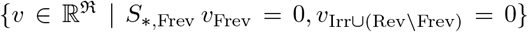. From 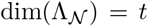, we get *ρ*(*S*_∗,Frev_) = |Frev| − *t*.

By Prop. 3.1, it follows that 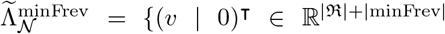 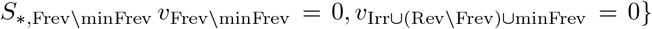. Since 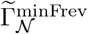 is pointed, this implies *ρ*(*S*_∗,Frev\minFrev_) = | Frev | − | minFrev | = | Frev | − *t* = *ρ*(*S*_∗,Frev_). For each *i*_*k*_ ∈ minFrev, the column 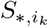 is therefore linearly dependent on the columns of *S*_∗,Frev\minFrev_. Thus, there exists a flux vector 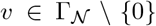 with 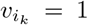 and supp(*v*) ⊆ (Frev \ minFrev) ∪ {*i*_*k*_}. It follows that 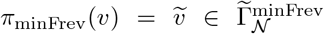 and 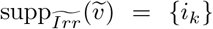, with 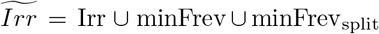. This implies that 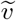 has minimal support w.r.t irreversible reactions in 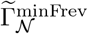 According to Prop. 2.2, 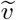 is an ER of 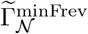.

Suppose there exists an ER 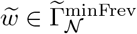 with 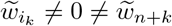. This would imply that 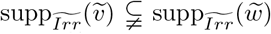, which is a contradiction. It follows that the set of ERs of 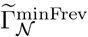 does not contain any 2-cycle.

From the proof of Prop. 5.1 we can deduce the following corollaries:

### Corollary 5.2

*Let* 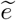 *be an extreme ray in* 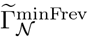. *Then there is at most one reaction j* ∈ minFrev ⋃ minFrev_split_ *with* 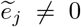. *If* 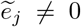, *for some j* ∈ minFrev ⋃ minFrev_split_, *then* 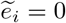, *for all i* ∈ Irr. *Moreover*, 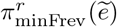 *is a reversible EFM of the original network* 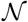.

### Corollary 5.3

For each *i*_*k*_ ∈ minFrev *there exist exactly two extreme rays* 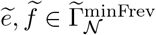 *such that* 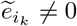 *and* 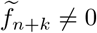. *After recombination, both* 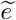 *and* 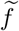 *define a single reversible EFM e in* 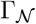 *with* 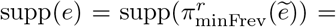 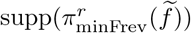. *Altogether, the t reactions in minFrev define* 2*t extreme rays in* 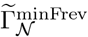 *and t linearly independent reversible EFMs in* 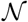.

The following proposition relates the supports of the extreme rays of 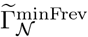 to the minimal metabolic behaviors (MMBs) of 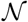.

### Proposition 5.4

*Let* minFrev *be a minimum set of fully reversible reactions in a metabolic network* 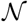 *such that* 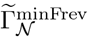 *is pointed. The non-empty supports* 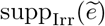 *of the extreme rays* 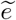 *of* 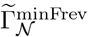 *are exactly the minimal metabolic behaviors of* 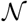.

**Proof:** Let *D* be an MMB in 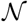 and let 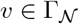 such that supp_Irr_(*v*) = *D*. By Lemma 2 in [18], there is a decomposition *v* = *e*+*y*, with *e*, 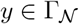, *e*_minFrev_ = 0 and *y*_Irr_ = 0. For 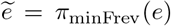 and 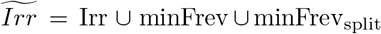 it follows that 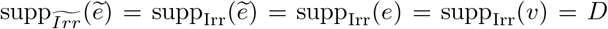. Suppose 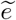 is not an extreme ray in 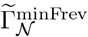. By Prop. 2.2, there exists 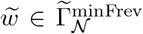 with 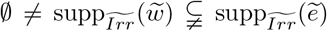. For 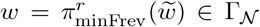 this implies 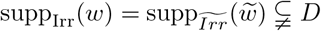, in contradiction to *D* being an MMB in Γ_*N*_.

Conversely, let 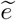 be an extreme ray in 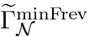 with 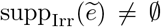. By Cor. 5.2, we have 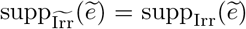. Assume 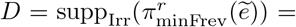 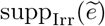 is not an MMB in 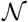. Then there exists 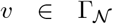 with 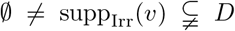. As before, we consider a decomposition *v* = *u* + *y*, with *u*, 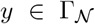, *u*_minFrev_ = 0 and *y*_Irr_ = 0. For 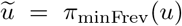 we get 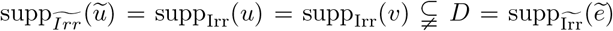, in contradiction to 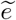 being an extreme ray in 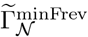.

We prove now one of our main theorems, stating that for any minimum set of fully reversible reactions minFrev, the extreme rays of 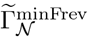 after recombination define a MEMo of 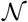.

### Theorem 5.5

*Let* minFrev *be a minimum set of fully reversible reactions in a metabolic network* 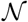 *such that* 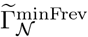 *is pointed. After recombination, the extreme rays of the pointed cone* 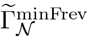 *define a minimum set of EFMs or MEMo in* 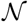. *Any set of EFMs of smaller cardinality cannot be a generating set for the ux cone* 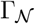.

**Proof:** According to [41, Sect. 8.8], a minimum set of generating vectors for 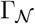 always consists of one vector in each minimal proper face of 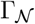 and a vector space basis of 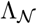. According to Cor. 5.2, there are two types of extreme rays 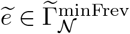:

Case 1: 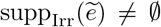 and 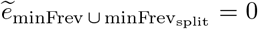. Let 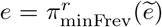 be the recombined vector. By Prop. 5.4, 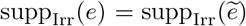 is an MMB *D* in 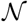. By [26] this implies that *e* belongs to the minimal proper face defined by *D*. Since supp_Irr_(*e*) ≠ 0, we also have 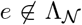. By Prop. 2.2, the extreme ray 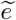 is an EFM in 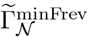. Thus, 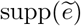 is inclusion-minimal in 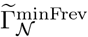. Since 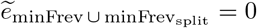, we have 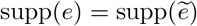, which implies that *e* is an EFM in 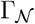 (otherwise, there would exist 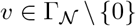 with 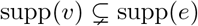, and for 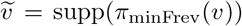) we would get 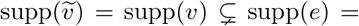 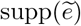, which is a contradiction).

Case 2: 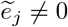 for some *j* ∈ minFrev ∪ minFrev_split_ and 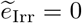. According to Cor. 5.2, all ERs with this property lead to *t* reversible EFMs in 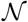, which after recombination generate 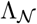.

In conclusion, after recombination, the ERs with the first property represent the MPFs of 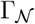 and the ERs with the second property form a basis of 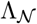. Altogether, the recombined ERs of 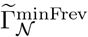 form a minimum set of EFMs generating 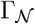, and therefore define a MEMo.

### Corollary 5.6

*For a given metabolic network* 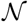 *the number of EFMs in a MEMo is always s* + *t, where s is the number of minimal proper faces in* 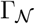 *and t is the dimension of* 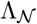.

Alg. 2 describes how to compute a MEMo given a minimum set of fully reversible reactions minFrev, which can be obtained by Alg. 1. First, the reactions in minFrev are split in order to get the pointed augmented flux cone 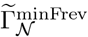. Next, we compute the ERs of 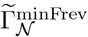 using an existing tool based on the double description method [12], such as `polco` [50] or `cdd` [11].

### Algorithm 2: Finding MEMo

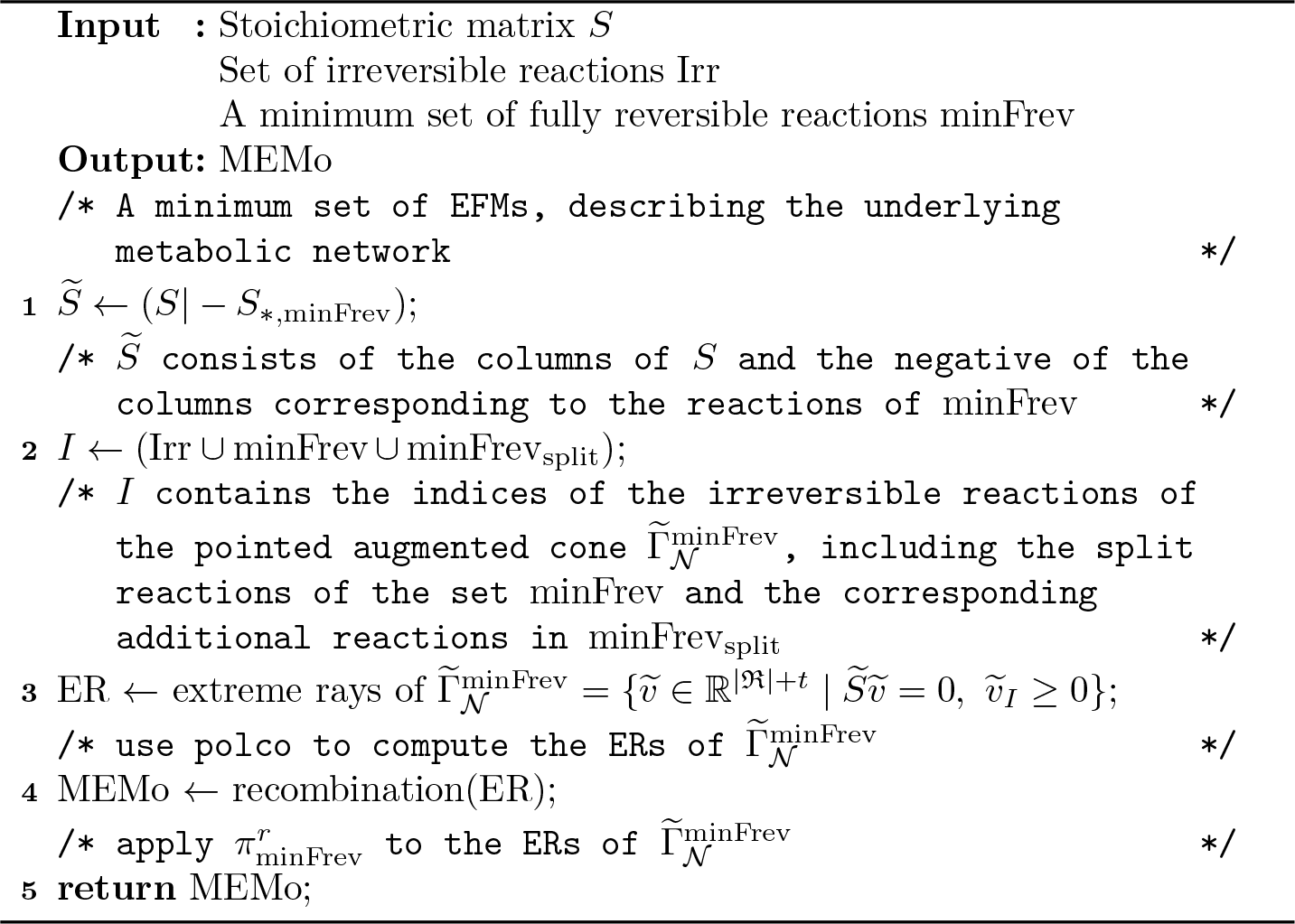

## 6 Computational results and discussion

In this section we first give some computational results, followed by a discussion and potential applications for MEMos.

### 6.1 Computational results

We implemented our method in `MATLAB` using `polco` [50] to compute the extreme rays of a pointed cone. Our software is available at https://sourceforge.net/projects/findingmemo. In the following, we present computational results for various metabolic network reconstructions taken from `BiGG` Models [22], `KEGG` [21] (together with `KEGGtranslator` [55]), and the `BioModels` Database [27]. Tab. 1 summarises the main characteristics of the metabolic networks studied. In Tab. 2, we study the size of the MEMos compared to the number of extreme pathways (EPs) and the number of EFMs, which were computed with `polco` [50] resp. `efmtool` [49]. In Fig. 9, we compare the number of flux vectors in a MEMo to the number of EPs and EFMs for those networks for which we were able to compute the EPs and EFMs. Since the number of EFMs of a MEMo is much smaller than the number of all EFMs, it is faster to compute them. A table with running times for all computations can be found in the Appendix.

**Figure 9.**
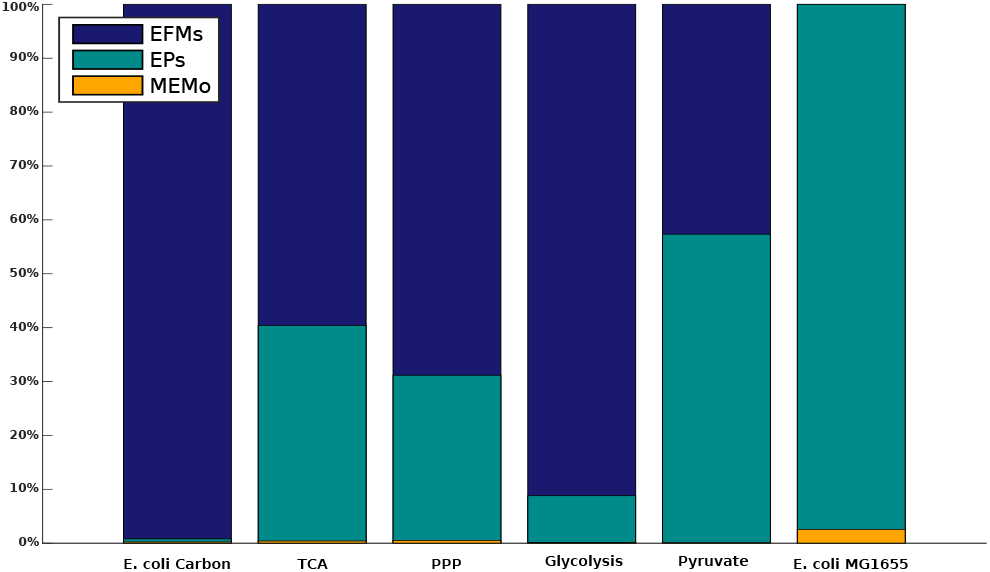
Comparison of the number of flux vectors of the three different generating sets we discussed in this article. We set the number of all EFMs to 100% and compare the vectors of the other sets accordingly. The bars overlay and the largest number is always on the top. Hence, if there is no blue bar for the EFMs *Escherichia coli* MG1655 the number of EPs and the number of EFMs are the same. Note that there is always a yellow bar indicating the size of the MEMo even it is hard to see. This shows how small the number of flux vectors in a MEMo is, compared to the whole set of EFMs.

**Table 1.** Characteristics of different metabolic networks used for the comparison in Tab. 2 **network**: name of the network. **rxns**: number of unblocked reactions. **mets**: number of metabolites. **rev**: number of unblocked reversible reactions. **intrev**: number of unblocked internal reversible reactions. **frev**: number of unblocked fully reversible reactions. **t**: dimension of the lineality space of the flux cone.

| **network** | **rxns** | **met**s | **rev** | **intrev** | **frev** | **t** |
| --- | --- | --- | --- | --- | --- | --- |
| *Escherichia coli* carbon metabolism [4] | 34 | 18 | 34 | 18 | 32 | 15 |
| Citrate cycle (TCA) [21] | 35 | 22 | 31 | 25 | 30 | 12 |
| Pentose phosphate pathway [21, 55] | 57 | 34 | 31 | 18 | 27 | 8 |
| Glycolysis / Gluconeogenesis [21, 55] | 61 | 32 | 42 | 28 | 33 | 13 |
| Pyruvate metabolism [21, 55] | 81 | 28 | 34 | 29 | 24 | 12 |
| *Escherichia coli* MG1655 [22] | 87 | 68 | 34 | 28 | 0 | 0 |
| *Rhizobium etli* iOR363 [27, 34] | 194 | 371 | 71 | 60 | 0 | 0 |
| *Buchnera* iSM197 [27, 29] | 261 | 255 | 17 | 11 | 0 | 0 |
| *Mycoplasma pneumoniae* iJW145 [27, 54] | 306 | 266 | 73 | 47 | 0 | 0 |
| *Blattabacterium cuenoti* iCG238 [15, 27] | 350 | 364 | 73 | 60 | 5 | 2 |
| *Mycoplasma genitalium* iPS189 [27, 47] | 351 | 346 | 107 | 26 | 3 | 1 |
| *Mannheimia succiniciproducens* iSH335 [16, 27] | 374 | 316 | 58 | 49 | 3 | 1 |
| *Blattabacterium* iCG230 [15, 27] | 412 | 358 | 71 | 59 | 5 | 2 |
| *Helicobacter pylori* 26695 [39] | 452 | 396 | 94 | 82 | 31 | 6 |
| *Human Erythrocyte* iAB-RBC-283 [2, 27] | 469 | 342 | 123 | 68 | 10 | 2 |
| *Helicobacter pylori* iCS291 [27, 39] | 493 | 396 | 94 | 67 | 31 | 6 |
| *Clostridium thermocellum* iSR432 [27, 36] | 581 | 564 | 95 | 95 | 32 | 20 |

**Table 2.** Size of MEMos, number of EPs and number of EFMs for different networks. **t**: dimension of the lineality space of the flux cone. **rxns**: number of unblocked reactions of the network. **network**: name of the network. **MEMo**: size of a minimum set of elementary modes. **EPs**: number of extreme pathways. **EFMs**: number of elementary flux modes. **N/A**: The programs used (`polco` [50] and `efmtool` [49]) run out of memory.

| **network** | **rxns** | **t** | **MEMo** | **EPs** | **EFMs** |
| --- | --- | --- | --- | --- | --- |
| *Escherichia coli carbon metabolism* [4] | 34 | 15 | 15 | 52 | 6,421 |
| Citrate cycle (TCA) [21] | 35 | 12 | 16 | 1,564 | 3,870 |
| Pentose phosphate pathway [21] | 57 | 8 | 27 | 1,607 | 5,155 |
| Glycolysis / Gluconeogenesis [21] | 61 | 13 | 29 | 1,716 | 19,464 |
| Pyruvate metabolism [21] | 81 | 12 | 49 | 27,361 | 47,708 |
| *Escherichia coli* MG1655 [22] | 87 | 0 | 2,572 | 100,274 | 100,274 |
| *Rhizobium etli* iOR363 [27, 34] | 194 | 0 | 6,147 | N/A | N/A |
| *Buchnera* iSM197 [27, 29] | 261 | 0 | 245 | 1,863 | N/A |
| *Mycoplasma pneumoniae* iJW145 [27, 54] | 306 | 0 | 1,593,880 | N/A | N/A |
| *Blattabacterium cuenoti* iCG238 [15] | 350 | 2 | 376 | 1,863 | N/A |
| *Mycoplasma genitalium* iPS189 [27, 47] | 351 | 1 | 190,471 | N/A | N/A |
| *Mannheimia succiniciproducens* iSH335 [16, 27] | 374 | 1 | 2,064,760 | N/A | N/A |
| *Blattabacterium* iCG230 [15, 27] | 412 | 2 | 84 | N/A | N/A |
| *Helicobacter pylori* 26695 [39] | 452 | 6 | 150,138 | N/A | N/A |
| *Human Erythrocyte* iAB-RBC-283 [2, 27] | 469 | 2 | 6,179 | N/A | N/A |
| *Helicobacter pylori* iCS291 [27, 39] | 493 | 6 | 150,138 | N/A | N/A |
| *Clostridium thermocellum* iSR432 [27, 36] | 581 | 20 | 1,039,267 | N/A | N/A |

In all networks that we considered the number of fully reversible reactions and the number of internal reversible reactions is greater than or equal to the dimension *t* of the lineality space 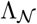. While there always exist internal reversible reactions, it can happen that there are no fully reversible reactions. In this case, *t* = 0 and the original flux cone is already pointed, i.e., there is no need to split any reaction before computing a MEMo. This holds for the networks *Escherichia coli* MG1655 [22], *Rhizobium etli* iOR363 [27, 34], *Buchnera* iSM197 [27,29], and *Mycoplasma pneumoniae* iJW145 [27,54]. For all these networks, there exist internal reversible reactions which are split when computing the extreme pathways (EPs). This is why the number of EPs is larger than the number of flux vectors in a MEMo. For example, instead of 2,572 EFMs in a MEMo for the network *Escherichia coli* MG1655 [22], there exist 100,274 EPs. For this network, the number of EFMs and EPs is the same, although not all reversible reactions are internal.

For the networks *Mannheimia succiniciproducens* iSH335 [16, 27], *Helicobacter pylori* 26695 [39], *Helicobacter pylori* iCS291 [27, 39], and *Clostridium thermocellum* iSR432 [27, 36] we were not able to compute the EPs because `polco` was running out of memory, while we were always able to compute a MEMo.

## 6.2 Discussion

When computing only a subset of EFMs and not the whole set, several issues arise. One drawback of computing a MEMo is that the resulting set of EFMs is not unique. This problem was already addressed by [44]. We prove in Sect. 5 that there exists a bijection between the irreversible EFMs in a MEMo and the MMBs of the network. Thus, the set of active irreversible reactions is the same for all flux vectors in all different MEMos, while the set of active reversible reactions is not, therefore giving rise to the non-uniqueness of MEMos.

The MEMos that can be obtained by Algor. 2 depend on the set minFrev of reactions that are split. As the following example shows, different minimum sets minFrev of fully reversible reactions may result in the same MEMo. Furthermore, there exist MEMos that cannot be obtained from any set minFrev.

### Example 6.1 (Different MEMos)

*For the network in Fig. 1 there exist three different reversible EFMs with the supports {3, 5, 9, 10}, {3, 5, 11, 12}, {9, 10, 11, 12}. Each two of them form a basis of the lineality space 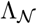, thus there are three bases. Furthermore, there exist three MMBs {2}, {6, 7}, {6, 8}, see Example 2.1, which correspond to three minimal proper faces of the flux cone 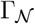. The minimal proper face for the MMB {6, 7} contains only one EFM with support {4, 6, 7}. The other two minimal proper faces each contain three EFMs, namely {1, 2, 3, 4}, {1, 2, 4, 5, 9, 10}, {1, 2, 4, 5, 11, 12} for the MMB {2} and {4, 6, 8, 9, 10}, {3, 4, 5, 6, 8}, {4, 6, 8, 11, 12} for the MMB {6, 8}. The remaining eight EFMs lie in the interior of the cone. These EFMs contain more than one MMB, e.g.{1, 2, 3, 6, 7} contains the MMBs {1, 2} and {6, 7}. Therefore these EFMs do not lie in a minimal proper face but in the interior of the cone. Thus they are not contained in any MEMo. By choosing one basis for the lineality space and one EFM for each minimal proper face, we get in total* 3 · 1· 3 · 3 = 27 *possible MEMos*.

*By enumeration, we can determine that there exist 12 different minimum sets of fully reversible reactions* minFrev *for the network in Fig. 1, which can be found in the Appendix. These 12 sets give rise to 5 different MEMos. For example, the two* minFrev *sets* {3, 9} *and* {3, 10} *result in the same MEMo. A MEMo that cannot be obtained by choosing a* minFrev *set and computing the extreme rays of the cone 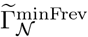 is the set of EFMs with the supports* {1, 2, 3, 4}, {4, 6, 7}, {4, 6, 8, 9, 10}, {3, 5, 9, 10}, {3, 5, 11, 12}.

Alg. 1 computes always the same set minFrev for a given network. Therefore, the MEMo which Alg. 2 computes, will be always the same too, although there exist different MEMos for a given network. Unfortunately, Alg. 2 cannot be used to compute all MEMos of a network even if all sets of minFrev are given, see Example 6.1.

Another issue, besides non-uniqueness, when studying only a subset of EFMs is that not all EFMs of biological relevance may be contained in a MEMo, as pointed out in [44]. This is a valid argument for computing the set of all EFMs. However, for genome-scale metabolic networks this is often not feasible, or the number of EFMs is too large to identify relevant EFMs. In such cases, a MEMo is a good alternative for studying the underlying metabolic network because it is smaller but still represents the whole network. As described in the following subsection, several methods exist which can use a generating subset of EFMs or MEMo instead of the whole set.

### 6.3 Applications

A typical application for EFMs is the decomposition of a given flux vector into a set of EFMs [45, 46]. Since the set of all EFMs is often not computable, different methods exist which use only a subset of EFMs [3, 17]. [53] introduced a method which decomposes flux vectors to compute the *α-spectrum* of a given flux vector using a (generating) subset of EFMs. The *α*-spectrum defines which EFMs can and cannot be included in the reconstruction of a given flux vector and to what extent they individually contribute to the decomposition. So far, either the set of all EFMs is used or the set of EPs. As shown in Tab. 2, the number of EPs may be much larger than the number of flux vectors in a MEMo. Furthermore, there exist networks where it is possible to compute a MEMo but not the EPs. For these networks, the *α*-spectrum can be computed using a MEMo, whereas using the set of EPs or the set of all EFMs is not possible. Based on Wiback’s work, [28] developed a method which computes the *α*-spectrum for a given flux vector including uncertainties. Their concept of *flux-spectrum* is used in scenarios where measurements are incomplete and/or uncertain. Since the method is based on the *α*-spectrum, again a generating subset or MEMo can be utilized instead of the whole set of EFMs.

Another well-known application of EFMs is the computation of *Minimal Cut Sets* (MCSs). MCSs are, like EFMs, minimal sets of reactions. Whenever the set of reactions of an MCS is removed from a metabolic network, certain predefined target behaviours are not possible anymore [24]. Given the whole set of EFMs, MCSs are inclusion-minimal hitting sets, i.e., sets of reactions such that each EFM contains at least one reaction of the hitting set [23]. Unfortunately, an arbitrary generating subset of EFMs cannot be used to compute MCSs. However, as shown in [38], one can use MMBs instead of EFMs to compute MCSs consisting only of irreversible reactions. Here, minimal hitting sets within the MMBs are enumerated, which correspond to *irreversible MCSs* (iMCSs). As shown in Sect. 5, there exists a bijection between the irreversible EFMs in a MEMo and the MMBs of the network. Given a MEMo, extracting the irreversible reactions from the EFMs in the MEMo provides the set of MMBs. In other words, when computing a MEMo we obtain at the same time the set of all MMBs. Therefore, the method presented here also gives a new way for computing MMBs, and based on these for computing iMCSs.

The EFMs in a MEMo can provide new insight into the importance of certain pathways or reactions in the network, especially since their irreversible part is unique. Indeed, investigating which reactions and EFMs are exchangeable and why, is an interesting task. Doing so, it would be possible to study the robustness of the underlying metabolic network. In addition, the EFMs of a MEMo could also be used to discover similarities to other networks by detecting subsets of EFMs that can be found in those other networks as well.

## 7 Related work

In the following, we discuss a number of papers that are closely related to our work.

### 7.1 Extreme pathways

In our approach, we compute a MEMo by splitting a minimum subset of reversible reactions. A closely related concept is the set of *extreme pathways* (EPs) [31, 40], which are computed by splitting all the internal reversible reactions. As shown in [31, 40], splitting only the *internal* reversible reactions always delivers an augmented flux cone which is pointed (given that there is only one exchange reaction per internal metabolite). This cone is unique and so are its extreme rays. After recombination, these extreme rays are called extreme pathways. The assumption that there exists only one reversible exchange reaction per metabolite is not true for all metabolic networks. Examples where this condition does not hold are the *Escherichia coli* carbon metabolism [4] or RECON1 [7, 22]. For computing a MEMo we do not need any condition on the network. However, the main advantage of our approach is that the number of EFMs in a MEMo is typically much smaller than the number of EPs, see Tab. 2.

### 7.2 Minimal metabolic behaviours

[26] introduce a unique minimum outer description of the flux cone 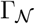 based on *minimal metabolic behaviours* (MMBs) and the lineality space 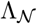. Each MMB corresponds to a minimal proper face of the flux cone or to set of flux vectors with the same minimum set of active irreversible reactions. Larhlimi and Bockmayr show that for each MMB there exists a corresponding EFM, but they do not consider reaction splitting and do not present a method for computing a minimal generating set consisting of EFMs only.

### 7.3 Minimal generating set

[18] introduce a method for finding a minimum generating set of flux vectors which corresponds to a set of EFMs and therefore is a MEMo. However, their approach is different from the one introduced here. We exploit splitting reversible reactions, while they use a method based on generating a basis of the null space of 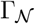. Additionally, they only prove that the reversible generating flux vectors are EFMs, but do not consider the irreversible flux vectors. We prove that all generating flux vectors obtained by our approach are EFMs. [18] only mention that the minimum set of EFMs is not unique. Here we discuss this issue in a systematic way and show how to obtain different MEMo’s by splitting different sets of reversible reactions. Furthermore, we provide a `MATLAB` tool to compute MEMos and illustrate the impact of our method by computational results for various genome-scale metabolic networks.

## 8 Conclusion and further work

In this paper we introduced the concept of a minimum set of elementary modes (MEMo) necessary to generate the flux cone of a metabolic network. We presented a method to compute these MEMos. We implemented our algorithm using `MATLAB` and showed that the size of MEMos often is by several orders of magnitude smaller than the number of EPs or EFMs. One drawback of the MEMos is that they are not unique because they depend on the set of fully reversible reactions which have to be split. However, one can show that the set of irreversible reactions involved in the MEMos is unique and that they will always be part of the MEMos independently of the reactions that are split. The biological relevance of the MEMos has to be further investigated. The need of having smaller sets of generating flux vectors has been addressed by various papers [1,9,18,20,35,40]. We expect that MEMos give rise to new opportunities for analysing large genome-scale metabolic networks.

## Acknowledgements

We thank Alexandra and Arne Reimers for fruitful discussions and comments on this paper.

## 9 Appendix

## 9.1 Example: all minFrevs and the corresponding MEMo

**Table 3.** All different sets of minFrev for the example network. The reactions of the given set minFrev are split and the extreme rays of the corresponding pointed flux cone are computed. After recombination the extreme rays always correspond to five different EFMs which form a MEMo. For different sets of minFrev the same set of EFMs can occur. **minFrev**: a set of two reactions. The augmented flux cone, after splitting these two reactions, is pointed. **EFMi**: the support of the *i*-th EFM of the MEMo. **MEMo**: which MEMo is described by the five EFMs. To point out which sets of minFrev result in the same MEMo we put a dividing line between different MEMos.

| minFrev | EFM1 | EFM2 | EFM3 | EFM4 | EFM5 | MEMo |
| --- | --- | --- | --- | --- | --- | --- |
| 3,9 | 1,2,4,5,11,12 | 3,5,11,12 | 4,6,7 | 4,6,8,11,12 | 9,10,11,12 | I |
| 3,10 | 1,2,4,5,11,12 | 3,5,11,12 | 4,6,7 | 4,6,8,11,12 | 9,10,11,12 | I |
| 3,11 | 1,2,4,5,9,10 | 3,5,9,10 | 4,6,7 | 4,6,8,9,10 | 9,10,11,12 | II |
| 3,12 | 1,2,4,5,9,10 | 3,5,9,10 | 4,6,7 | 4,6,8,9,10 | 9,10,11,12 | II |
| 5,9 | 1,2,3,4 | 3,5,11,12 | 4,6,7 | 4,6,8,11,12 | 9,10,11,12 | III |
| 5,10 | 1,2,3,4 | 3,5,11,12 | 4,6,7 | 4,6,8,11,12 | 9,10,11,12 | III |
| 5,11 | 1,2,3,4 | 3,5,9,10 | 4,6,7 | 4,6,8,9,10 | 9,10,11,12 | IV |
| 5,12 | 1,2,3,4 | 3,5,9,10 | 4,6,7 | 4,6,8,9,10 | 9,10,11,12 | IV |
| 9,11 | 1,2,3,4 | 3,4,5,6,8 | 3,5,9,10 | 3,5,11,12 | 4,6,7 | V |
| 9,12 | 1,2,3,4 | 3,4,5,6,8 | 3,5,9,10 | 3,5,11,12 | 4,6,7 | V |
| 10,11 | 1,2,3,4 | 3,4,5,6,8 | 3,5,9,10 | 3,5,11,12 | 4,6,7 | V |
| 10,12 | 1,2,3,4 | 3,4,5,6,8 | 3,5,9,10 | 3,5,11,12 | 4,6,7 | V |

## 9.2 Time for computation

**Table 4.** Time to compute a MEMo, EPs and EFMs for different networks. **rxns**: number of unblocked reactions of the network. **network**: name of the network. **MEMo**: time to compute a minimum set of elementary modes. **EPs**: time to compute the extreme pathways. **EFMs**: time to compute the elementary flux modes. **N/A**: The programs used (`polco` [50] and `efmtool` [49]) run out of memory.

| network | rxns | MEMo | EPs | EFMs |
| --- | --- | --- | --- | --- |
| *Escherichia coli carbon metabolism* [4] | 34 | 0.62 sec | 6 sec | 3 sec |
| Citrate cycle (TCA) [21] | 35 | 0.34 sec | 0.6 sec | 0.3 sec |
| Pentose phosphate pathway [21] | 57 | 0.5 sec | 0.4 sec | 0.5 sec |
| Glycolysis / Gluconeogenesis [21] | 61 | 0.5 sec | 0.3 sec | 0.9 sec |
| Pyruvate metabolism [21] | 81 | 0.5 sec | 1.4 sec | 2 sec |
| *Escherichia coli* MG1655 [22] | 87 | 1.6 sec | 17 sec | 9 sec |
| *Rhizobium etli* iOR363 [27,34] | 194 | 5 sec | N/A | N/A |
| *Buchnera* iSM197 [27,29] | 261 | 2.5 sec | 2 sec | N/A |
| *Mycoplasma pneumoniae* iJW145 [27,54] | 306 | 5 min | N/A | N/A |
| *Blattabacterium cuenoti* iCG238 [15] | 350 | 10.6 sec | 2 sec | N/A |
| *Mycoplasma genitalium* iPS189 [27,47] | 351 | 3 min | N/A | N/A |
| *Mannheimia succiniciproducens* iSH335 [16,27] | 374 | 3 min | N/A | N/A |
| *Blattabacterium* iCG230 [15,27] | 412 | 7.8 sec | N/A | N/A |
| *Helicobacter pylori* 26695 [39] | 452 | 1 min | N/A | N/A |
| *Human Erythrocyte* iAB-RBC-283 [2,27] | 469 | 15 sec | N/A | N/A |
| *Helicobacter pylori* iCS291 [27,39] | 493 | 1 min 5 sec | N/A | N/A |
| *Clostridium thermocellum* iSR432 [27,36] | 581 | 6.5 min | N/A | N/A |

